# The Role Of *Fzd8* For Bone Development And Homeostasis In A Mouse Model Generated By CRISPR/Cas9 Genome Editing

**DOI:** 10.1101/2025.01.19.633799

**Authors:** Zhengkun Lin, Jianquan He, Hui Huang, Xiaomei Lin, Heqing Chen, Wen Zhang, Jian Chen

**Affiliations:** Department of Rehabilitation, Zhongshan Hospital of Xiamen University, School of Medicine, Xiamen University, Xiamen, 361004, China; Icahn School of Medicine at Mount Sinai, New York, NY 10029, USA; Xiamen Humanity Rehabilitation Hospital, Xiamen, 361006, China

**Keywords:** osteoporosis, *FZD8*, mouse model, CRISPR/Cas9, differentially expressed genes

## Abstract

**Background:** *FZD8* could be a promising therapeutic target in osteoporosis (OP), although the signal transduction mechanism in OP regarding *FZD8* has not been completely elucidated.

**Aims:** We used the CRISPR/Cas9 technique to develop an *Fzd8*-knockout mouse model to study whether *Fzd8* inactivation results in genetic changes with potential correlations to OP.

**Materials and Methods:** Genotypes of distinguished classified knockout mice, i.e., heterozygous, homozygous, and wild-type were identified through PCR. Applying the murine model, third generation mice were used for the downstream experiments. We investigated the potential relevance of differentially expressed genes (DEGs) in OP.

**Results:** We found that osteoclasts significantly increased in *Fzd8*-knockout homozygous mice, compared to wild-type mice, while osteoblasts reduced significantly. Before transcription, heterozygous and homozygous mice possessed DEGs related to exons SNP, which are associated with exons CNV. After transcription, DEGs related to exons SNP in heterozygous and homozygous mice were observed, some of which are potentially associated with OP based on pathway and gene set enrichment analyses.

**Conclusions:** Our *Fzd8-*knockout murine model showed that there were significant alternations in *Fzd10* and *Lta* gene expressions and *Itgb3 and RANK* protein expressions among the wild-type and homozygous mice, which are significantly associated with bone remodeling. Our results revealed that *FZD8* could be a therapeutic target in OP. This study elucidates the molecular mechanisms in OP, providing evidence-based data for OP drug development and treatment.

## Introduction

Osteoporosis (OP) is a complicated metabolic bone disorder characterized by decreased bone mass, compromised bone microarchitecture, and increased fracture risk. The fractures and other complications caused by OP seriously affect the patient’s quality of life, leading to a heavy burden on their families, as well as on society. Consequently, OP has emerged as one of the most urgent issues in global public health [1].

Former investigations by the Osteoporosis Foundation [2] summarized that OP is prevalent in 13% of the Chinese population. It is speculated that the number of patients with OP or low bone mineral density in China will reach 212 million by 2050. Studies suggest that the canonical *Wnt* signaling pathway could enhance osteoclast (OC) differentiation and proliferation [3]. The non-canonical *Wnt* pathway affects bone remodeling as well. For instance, *Wnt16* could suppress OC generation, which prevents fractures due to fragility [4].

The frizzled (Fzd) protein is a transmembrane receptor responsible for binding to the Wnt protein, which activates intracellular signal transduction molecules [5]. Bone mineral contents and secreted frizzled-related proteins (sFRPs) can compete with *Fzd* receptors on the cell membrane surface to suppress *Wnt* protein function in bone pathophysiology [6]. *Fzd* receptors lack the low-density lipoprotein receptor-related protein 5 *(LRP5)* and low-density lipoprotein receptor-related protein 6 *(LRP6)*. Secreted form with respect to *FZD8,* cysteine-rich domains would antagonize *Wnt3a*-induced accumulation of *β*-catenin in mouse fibroblasts [7]. *Fzd8* is a *Wnt* receptor and is considered a *β*-catenin-independent pathway [8].

However, phenotype is not related to impaired *Wnt* signaling or osteoprotegerin *(OPG)* production through osteoblasts (OBs), which are irrelevant to bone formation [9]. Theoretical investigations have discovered that profiling multiple variants associated with bone phenotypes could improve fracture prediction accuracy [10]. *Fzd8* might be a therapeutic target for OP, though its signal transduction mechanism has not been clarified. In recent years, gene editing techniques have developed swiftly [11], such as the clustered regularly interspaced short palindromic repeat (CRISPR)/CRISPR-associated protein 9 (Cas9) technique, which can target specific genes of high interest [12]. There are few studies on the effect of *Fzd8*-knockout on OP, using whole genome sequencing (WGS). This study provides evidence-based medicine ideas for drug developments, which sheds new lights on OP rehabilitation diagnosis.

## Materials and methods

### Animals

Our lab housed mice with temperature-/humidity-controlled environment (23℃ ± 3℃/ 70% ± 10%) on a 12-h light/12-h dark cycle. Technicians maintained mice on standard mouse chow with 18.0% protein, 4.5% fat, 58% carbohydrate with water in Zhongshan Hospital Xiamen University. All mice were sacrificed when they aged between 20 and 30 weeks. We purchased C57BL/6J mice from the Animal Experiment Center of Fujian Province. The Fzd8 knockout mice (Zhongshan Hospital Xiamen University) were generated and maintained with C57BL/6J background. All animal experiments were carried out in Zhongshan Hospital Xiamen University and approved by Animal Subjects Committee in Zhongshan Hospital Xiamen University (approval No. XMVLAC20120044).

### *Fzd8**-***knockout mouse model

The *Fzd8*-knockout mouse model was designed and developed by the Shanghai Model Organisms Center, Inc. (Shanghai, China). The Cas9 mRNA was transcribed *in vitro* using the mMESSAGE mMACHINE T7 Ultra Kit (Ambion, TX, USA), following standard procedures (**Supplementary Figure S1**). The 4 single guide RNAs (sgRNAs) targeted to delete the exon 3 of the *Fzd8* gene were: 5’-AGGGAGTGGATCTCAAGCCTTGG-3’; 5’-TTACTCTGGGAGGTAGGGAGTGG-3’; 5’-CCGTAGTAAGAAGCTGAGTTAGG-3’; and, 5’-CCAAAGAGAAGGGTGCGGGCGGG-3’. We transcribed the sgRNAs *in vitro* using the MEGAshortscript Kit (Thermo Fisher Scientific, USA). We verified the first generation (F_0_) mice through PCR, using the following primer pairs F-5’-CGAACTCTTGGCAGGTCTGT-3’ and R-5’-ATGCCCATTGGAGCCATGAA-3’. We selected positive *Fzd8-*knockout F_0_ mice and mated them with the C57BL/6J mice to obtain the second generation (F_1_) heterozygous *Fzd8*-knockout mice. We intercrossed female and male F_1_ heterozygous mice to to produce F2 homozygous (HO) *Fzd8*-knockout mice.

### Micro-CT detection

We used the SkyScan 1176 (Bruker, Aartselaar, Belgium) system to measure different bone phenotype parameters. The NRecon software was used for 3D image reconstruction and viewing. The CTan software (version 1.13) was applied for bone analysis. We conducted micro-CT examinations of the left mice femora frozen at -40 ℃ in a freezer.

### Hematoxylin and Eosin (HE) staining

Mouse bones were fixed with 10% formalin and embedded in paraffin. Thereafter, slices in distilled water were dyed in a hematoxylin aqueous solution, which we then separated into acid and ammonia water. Slices were cleaned under running water for 1 h, and were then immersed in distilled water and dehydrated in 90% alcohol for 10 min. The size and morphology of the stained osteocytes in one section were investigated.

### Immunohistochemical (IHC) staining

Before dewaxing, we placed all tissue sections at room temperature (20-25℃) for 1 h, and then baked in a 60 ℃ thermostat for 20 min. The tissue sections were immersed in xylene for 10 min and soaked for another 10 min after altering xylene. The mouse bones were fixed in 10% formalin, embedded in paraffin, and sliced into 10 μm sections. The sections were mounted on slides, and antigen retrieval was conducted by incubating them with EDTA at 90 ℃ for 10 min. We then incubated the sections with 0.3% H_2_O_2_ for 0.5 h, followed by applying a blocking solution buffer for 1 h, at room temperature (20-25℃). We added 50 μL of the Anti-Wnt3a (Abcam; No. ab219412), Anti-RANK (Abcam; No. ab13918), and Anti-Integrin beta 3 (Abcam; No. ab179473) antibodies. We left the sections at room temperature (20-25℃) for 1 h, and cleaned them with PBS for 5 min, thrice. Next, we added 40-50 μL Goat Anti-Rabbit IgG H&L (Abcam; No. ab150077), and Goat Anti-Rabbit IgG H&L (Abcam; No. ab97051). The Image-Pro Expess analysis system (Informer Technologies, Inc 6.0) was utilized to quantify grayscale.

### Enzyme-linked immunosorbent assay (ELISA)

We placed the ELISA kit (Shanghai Pan Ke Industrial Co., LTD. No. 20200004) at room temperature (20-25℃), for 15-30 min. We erased the labeling plate to add 50 μL standard solution into the blank micropores. We added 10 μL biotin into the sample well, excluding the blank control well, and added 100 μL enzyme-labeled solution to each well. We sealed the enzyme label plate with a sealant and incubated it at 37 ℃ for 1 h. The absorbance (OD value) for each well was sequentially detected using a blank air conditioner at 450 nm wavelength.

### RNA-sequencing (RNA-seq)

Tail tissues of 8 wild-type (WT) and HO mice each, were separated into two groups for RNA-seq. Total RNA was obtained using the TRIzol reagent (Sigma), following standard protocols. RNA integrity was tested using the Bioanalyzer 2100 (Agilent). The cDNA libraries were checked for quality, and were further quantified using the 2100 Bioanalyzer. Each library was sequenced with Illumina Sequencing Kit (Illumina/TruSeq RNA Library Preparation Kit v2, /RS-122-2001/1 Ea) on one lane of NovaSeq 6000 sequencing system to obtain 150 bp paired-end reads.

### Whole genome sequencing (WGS)

Total genome DNA was extracted from mouse tails homogenized in 3 volumes of DNA extraction buffer using the Teflon homogenizer. The homogenized samples were incubated in a water bath at 55 ℃. Phenol and chloroform extraction, followed by isopropanol precipitation using 0.2× volume of 10 M ammonium acetate, was conducted at 7500 × g for 10 min. Two paired-end (PE) libraries were made for the WGS, which were sequenced on NovaSeq 6000 sequencing system (Illumina, San Diego, CA, USA). All sequencing-relevant procedures were carried out following protocols (Illumina, FC-121-4002 San Diego, CA, USA).

### Validation using PCR

PCR was used to verify the assembly features (**Supplementary Table S1**) developed using 1μL DNA template in each reaction and mixed with the Golden_Star_T6_Super_PCR_Mix (TSE101, Tsingke Biotechnology Co., Ltd.). PCR was run using the A300 PCR instrument (Hangzhou LongGene Scientific Instrument Co., LTD), applying a 2-step fast PCR with a 2 s denaturation step at 98 ℃, along with a 2 min annealing and extension step at 72 ℃ for 35 cycles.

### Validation using quantitative PCR (qPCR)

The assembly features developed were validated using qPCR (detailed information on qPCR primers is provided in **Supplementary Table S2**), using 5 μL RNA template in each reaction, mixed with RT6 cDNA Synthesis Kit Ver. 2 (Tsingke Biotechnology Co., Ltd. (TSK302S)). We conducted a 3-step fast qPCR (BIOER FQD-96A) using a 2 s denaturation step at 95 ℃. The cDNA developed was added to 10 μL ddH_2_O.

### Statistical evaluation criteria

All data analyses were performed using SPSS (version 22.0), and *P* < 0.05 was considered statistically significant. Each group of continuous variables was represented as mean ± standard deviation (SD). Median, interquartile range, and classified variable standard error were also calculated. Pearson’s Chi-square test or Fisher’s exact test were used for categorical variable analyses. Baseline characteristics and comparisons of major or minor results between the groups were also analyzed. To adjust confounding, a linear regression model was applied to the continuous variables.

### Bioinformatics data processing

We identified up-regulated and down-regulated genes in accordance with the log_2_ (*Fold Change*) and *P* < 0.05, indicating significance levels. Samples were clustered in terms of differentially expressed genes that were identified using the fragments per kilobase of exon per million mapped fragments. Correlations between samples were analyzed according to phenotypes. The Pearson correlation test was employed to obtain Pearson values regarding homogeneous mice. We performed gene ontology enrichment analysis; *q* values denoted significance, informing transcriptomic expressions of molecular processes. We performed gene set enrichment analysis, and the number of genes in each pathway pattern were analyzed. Standard procedures and pipelines were applied while performing these analyses[13].

## Results

### Baseline comparison between WT and HO mice

Statistical analyses suggested that there were no statistically significant differences in baseline indicators between the WT and HO mice, including their weights [mean ± standard deviation (SD): 22.40 ± 1.13 g (WT) vs. 21.70 ± 1.82 g (HO), P=0.403], blood glucose (*GLU*) (deviation: 6.20 ± 0.76 (WT) vs. 6.07 ± 1.41 (HO), *P*=0.749), N-terminal propeptide of type I procollagen (*P1NP)* (deviation: 5.69 ± 0.83 (WT) vs. 5.80 ± 1.36 (HO), *P*=0.857), and C-terminal cross-linking telopeptide *(CTX)* (deviation: 120.68 ± 22.44 (WT) vs. 109.42 ± 18.61 (HO), *P*=0.327) levels. The N-terminal propeptide of type I procollagen (*P1NP2)* levels in the HO mice were lower compared to WT mice, and the C-terminal cross-linking telopeptide *(CTX2)* levels in HO mice were higher (**Figure 1**).

**Figure 1.**
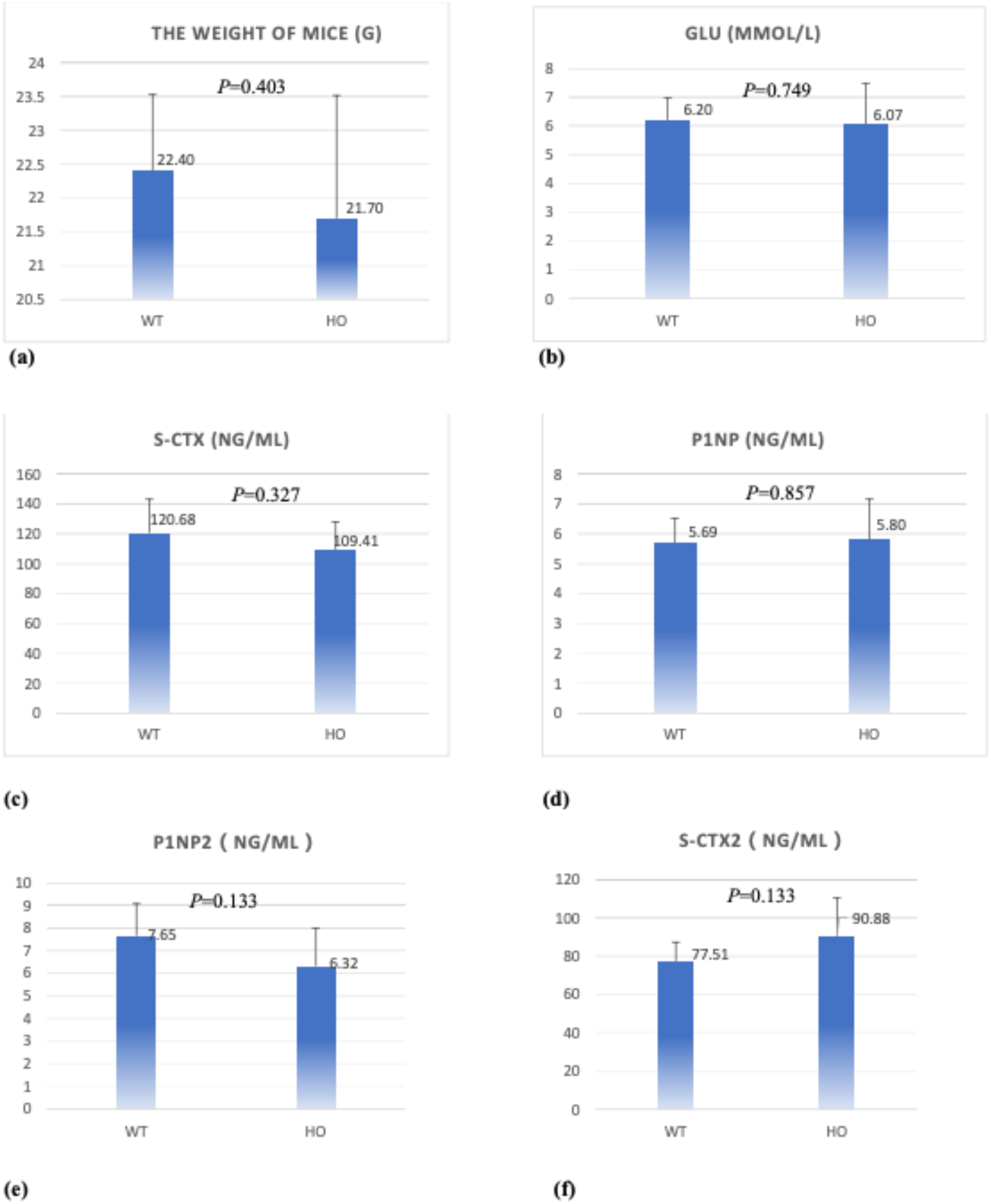
Comparison of WT and HO mice. The comparison of the weight **(a)** of the two groups of mice. The comparison of the blood glucose (GLU) **(b)**, N-terminal propeptide of type I procollagen (P1NP) **(c)**, and S-CTX **(d)** levels between the WT and HO mice before they euthanization. The comparison of the P1NP2 **(e)** and S-CTX2 **(f)** levels between the WT and HO mice after they were euthanized.

### Micro-CT detection

There were significant differences in the various bone imaging indices between the WT and HO groups (**Supplementary Figure S2**), such as the bone mineral density (BMD) of the cortical and trabecular bones, the total volume (TV), and bone volume (BV) of the cortical bone, the BV/TV of the cortical bone, among others. The mean and SD of bone phenotypic indicators in the WT and HO mice are provided in (**Supplementary Table S3**). Among these indicators, the mean of the phenotypic indicators in the HO mice is less than that in the WT mice. Trabeculae, BV, and BMD decreased after *FZD8*-knockout. The bone trabeculae, bone cortex, and BV in the HO mice reduced significantly after *Fzd8*-knockout (**Figure 2**).

**Figure 2.**
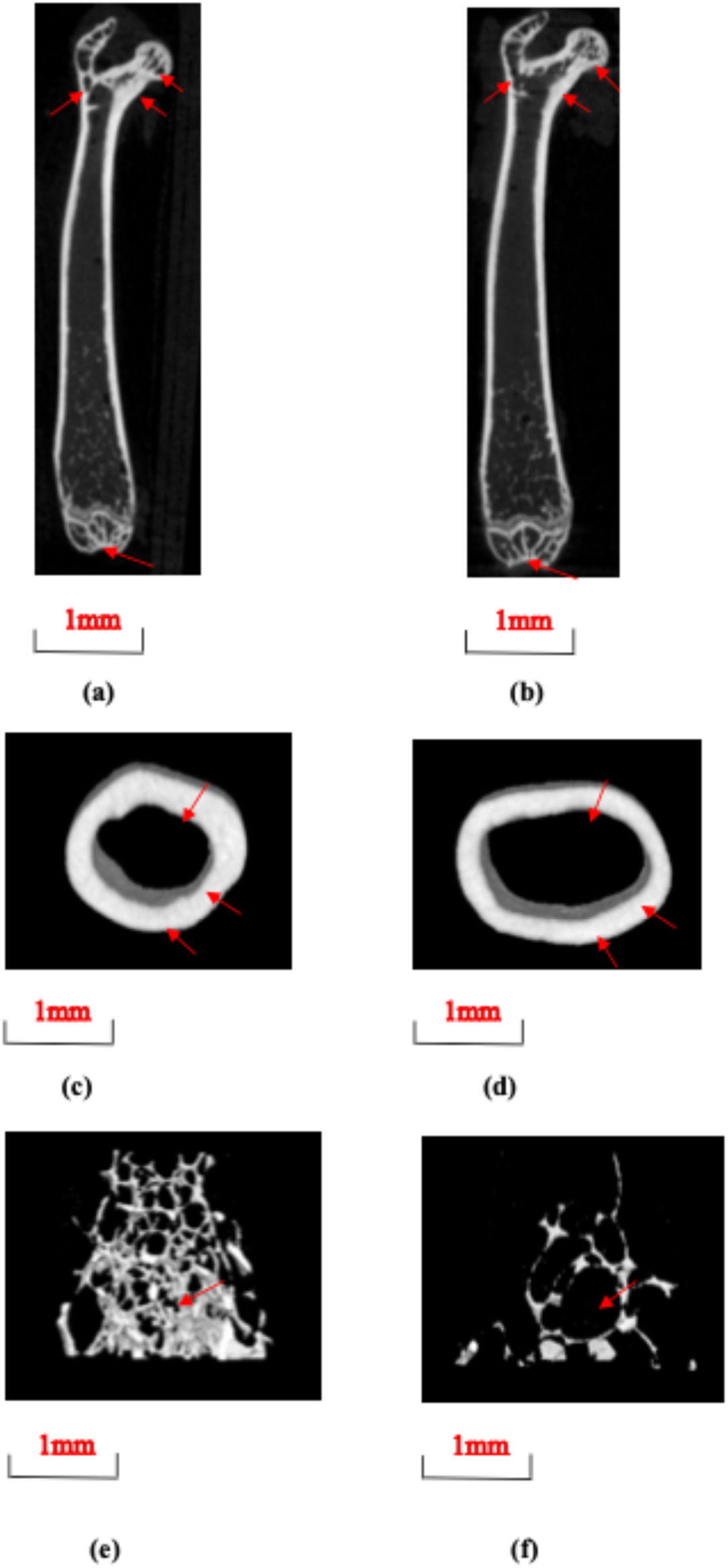
The coronal, cross-sectional, and 3D view of the middle and upper parts of the left femur in WT and HO mice. Coronal view of the middle and upper parts of the left femur in WT **(a)** and HO **(b)** mice. Cortical and trabecular bones of the HO mice have lower bone density, thinner cortices, and greater trabecular spacing, compared to the WT mice, as denoted by arrows. Cross-sectional view of the middle and upper parts of the left femur in WT **(c)** and HO **(d)** mice. Cortical bones of the HO mice have a lower bone density compared to the WT mice, as denoted by arrows. 3D view of the middle and upper parts of the left femur in WT **(e)** and HO **(f)** mice. Cortical and trabecular bones of the HO mice have lower bone density, thinner cortices, and greater trabecular spacing compared to the WT mice, as denoted by arrows.

### Differences in protein expression between WT and HO mice

There were significant differences among the *RANK*, *Wnt3a*, and *Itgb3* protein expressions in the WT and HO mice. Independent experienced pathologists evaluated the immunoreactivity of the *RANK*, *Wnt3a*, and *Itgb3* proteins in the WT and HO mice (**Figure 3-1**). *RANK*, *Wnt3a*, and *Itgb3* positive cells are represented by arrows. With the increase in age, various organ functions exhibited physiological degeneration and a pro-inflammatory state; identified by the low immune functions and decreased motor functions, among others. The HE staining charts (**Figure 3-1**) show that the HO mice have significantly fewer bone trabeculae than the WT mice.

**Figure 3-1.**
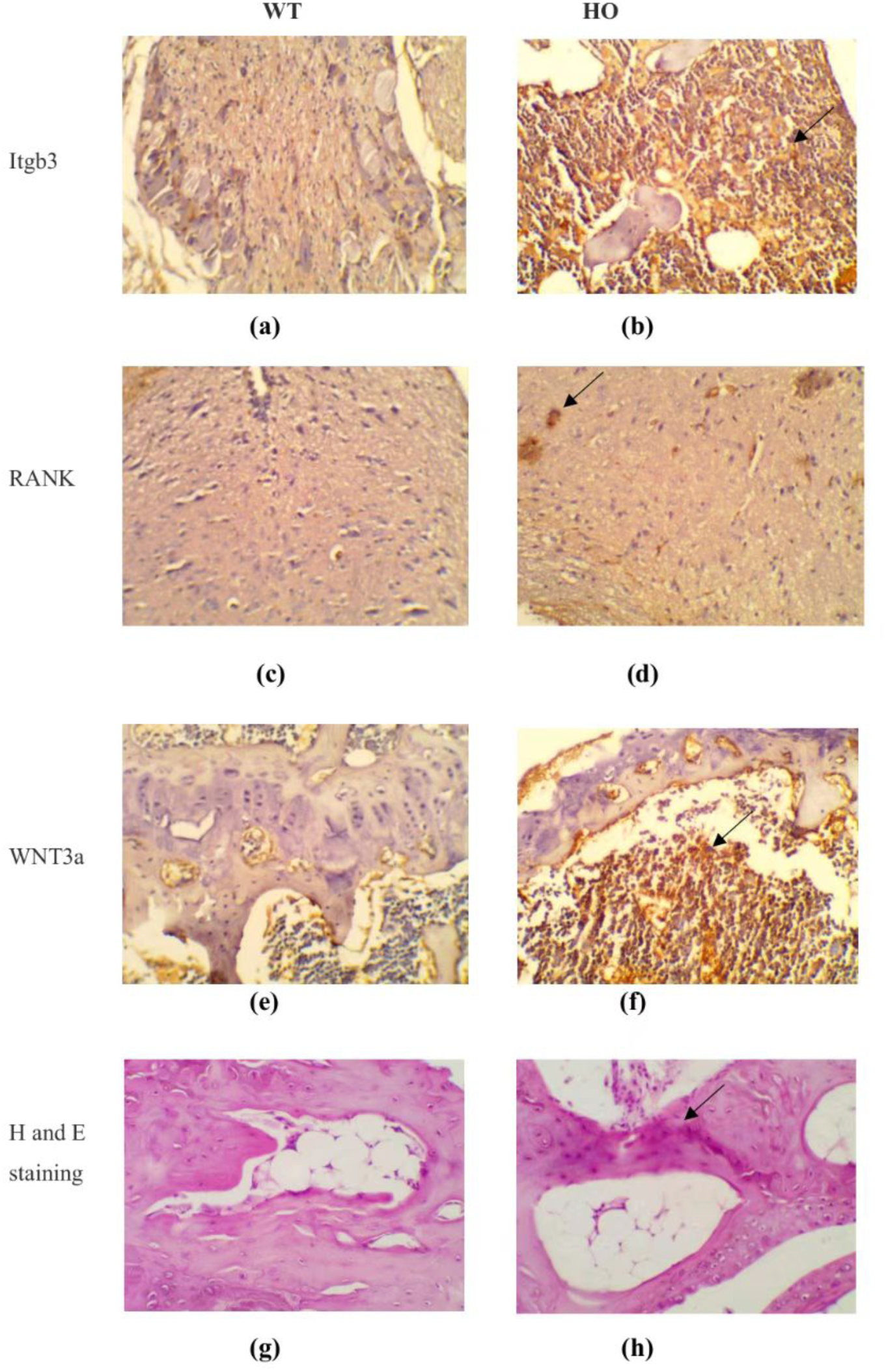
Immunoreactivity of the *Wnt3a*, *Itgb3*, and *RANK* proteins, along with HE staining. Independent experienced pathologists evaluated the immunoreactivity of the *RANK Wnt3a*, and *Itgb3* proteins, between the WT and HO mice. *RANK*, *Wnt3a*, and *Itgb3* positive cells were detected in bone tissues, as denoted by arrows (In immunohistochemistry, the positive staining is brownish yellow or brown particles). Representative images of H&E staining in the femoral metaphysis of different groups of mice (X200) are also provided. **(a-b)** *Itgb3* was expressed in WT and HO mice, and positive expression was confirmed by the brown-yellow color; *Itgb3* expression in HO mice was significantly higher than that in WT mice. **(c-d)** *RANK* was expressed in both WT and HO mice, and positive expression was confirmed by the brown-yellow color; *RANK* expression in HO mice was significantly higher than that in WT mice. **(e-f)** *Wnt3a* was expressed in both WT and HO mice, and positive expression was confirmed by the brown-yellow color; *Wnt3a* expression in HO mice was significantly higher than that in WT mice. **(g-h)** Under high magnification microscope (X200), the *Itgb3*, *Wnt3a*, and *RANK* protein expressions in HO mice were higher than that in WT mice.

All factors transmit information through the stimulated *OPG*/*RANK*/*RANKL* signal transduction system, which could reduce *OPG*/*RANKL* ratios and suppress OB function. *β*-catenin is the key factor in the *Wnt/β*-catenin signaling pathway. *RANK* and *Itgb3* expressions were different in the two mice groups (WT and HO) due to *Fzd8*-knockout (**Figure 3-2**). Combined with the influence of *RANK* on OP, *Fzd8-*knockout affects OP indirectly.

**Figure 3-2.**
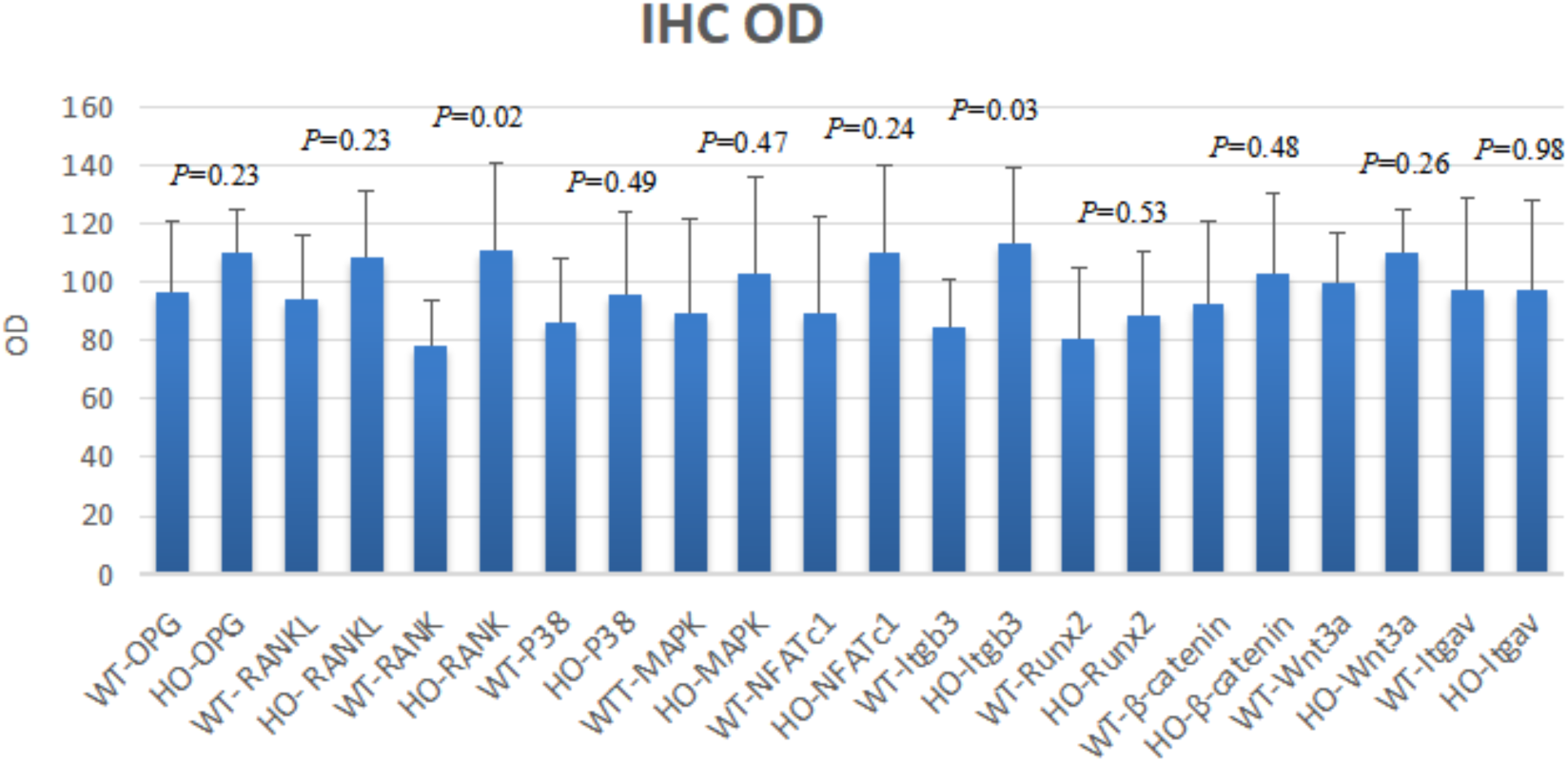
Comparison of the differences in immunohistochemical protein expressions between WT and HO mice after *Fzd8*-knockout. The *P* value represents the differences in the expression levels of each immunohistochemical protein in the WT and HO mice after *Fzd8* knockout; *P* < 0.05 indicates a statistically significant difference in the protein expression level between the two groups, and *P* > 0.05 indicates that the expression level of the immunohistochemical protein was not significantly different between the two groups.

### Gene expression differences between WT and HO mice

Expression levels of *Fzd10* and lipoteichoic acid (*Lta*) were statistically different in the WT and HO mice (**Figure 4**). *Fzd10* expression was significantly down-regulated, while *Lta* expression was noticeably up-regulated. In this study, several OP-related genes, including *Fzd10* in the canonical *Wnt* signaling pathway, are predicted to be the target genes of differentially expressed miRNAs, and *Fzd10* expression was down-regulated in OP.

**Figure 4.**
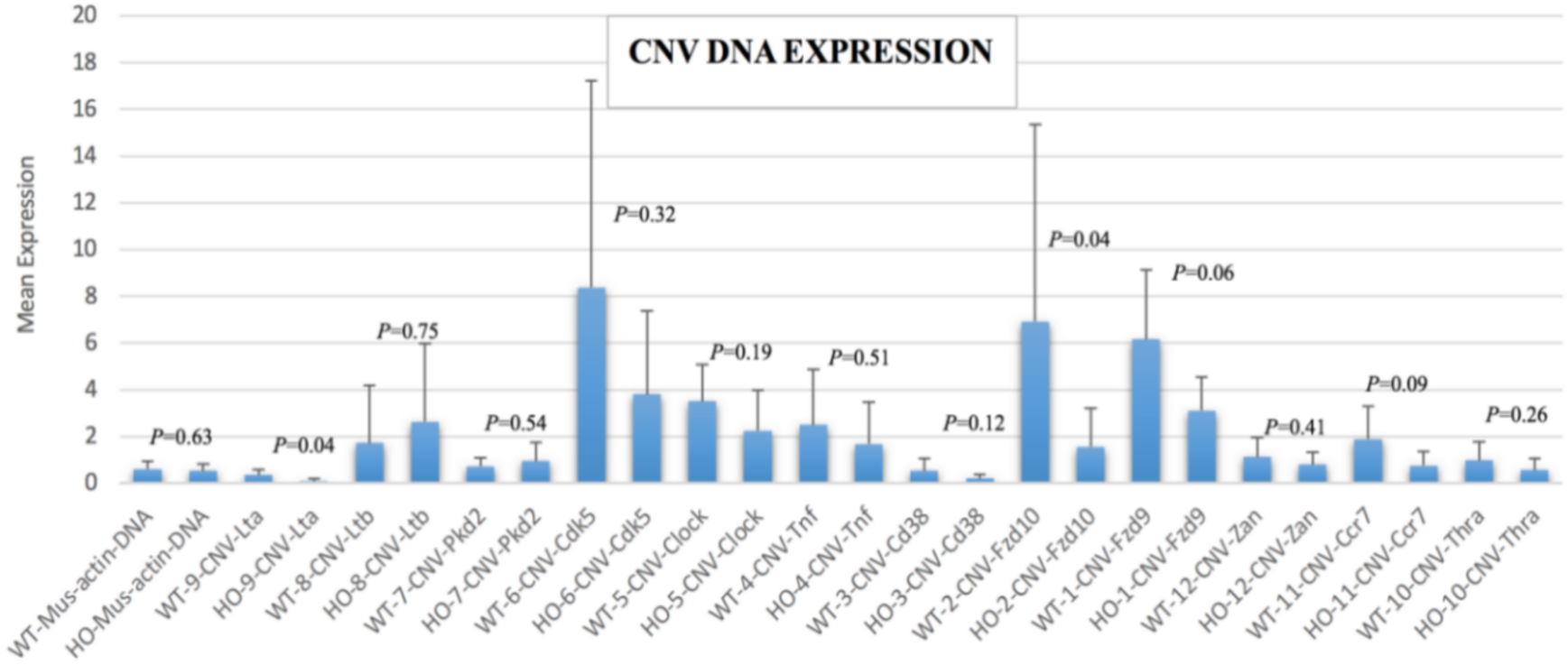
Comparison of differences in gene expression in WT and HO mice after *Fzd8*-knockout. The *P* value represents the difference between the expression levels of each gene in WT and HO mice after *Fzd8*-knockout; *P* < 0.05 indicates that the expression level of the gene was significantly different between the two groups, and *P* > 0.05 indicates that the expression level of the gene was not significantly different between the two groups.

### Bioinformatics analysis outputs

We found that the genes *Col1a1*, *Col1a2*, *Col3a1*, *Ibsp*, *S100a8*, *BC100530*, *Myoc*, *Col6a2*, and *Pcdh12* were significantly differentially expressed (*P* < 0.01; **Figure 5**). These genes are closely related to OP, further elucidating the effects of *Fzd8*-knockout in OP. *Col1a1* is the most significant gene that was up-regulated in this study.

**Figure 5.**
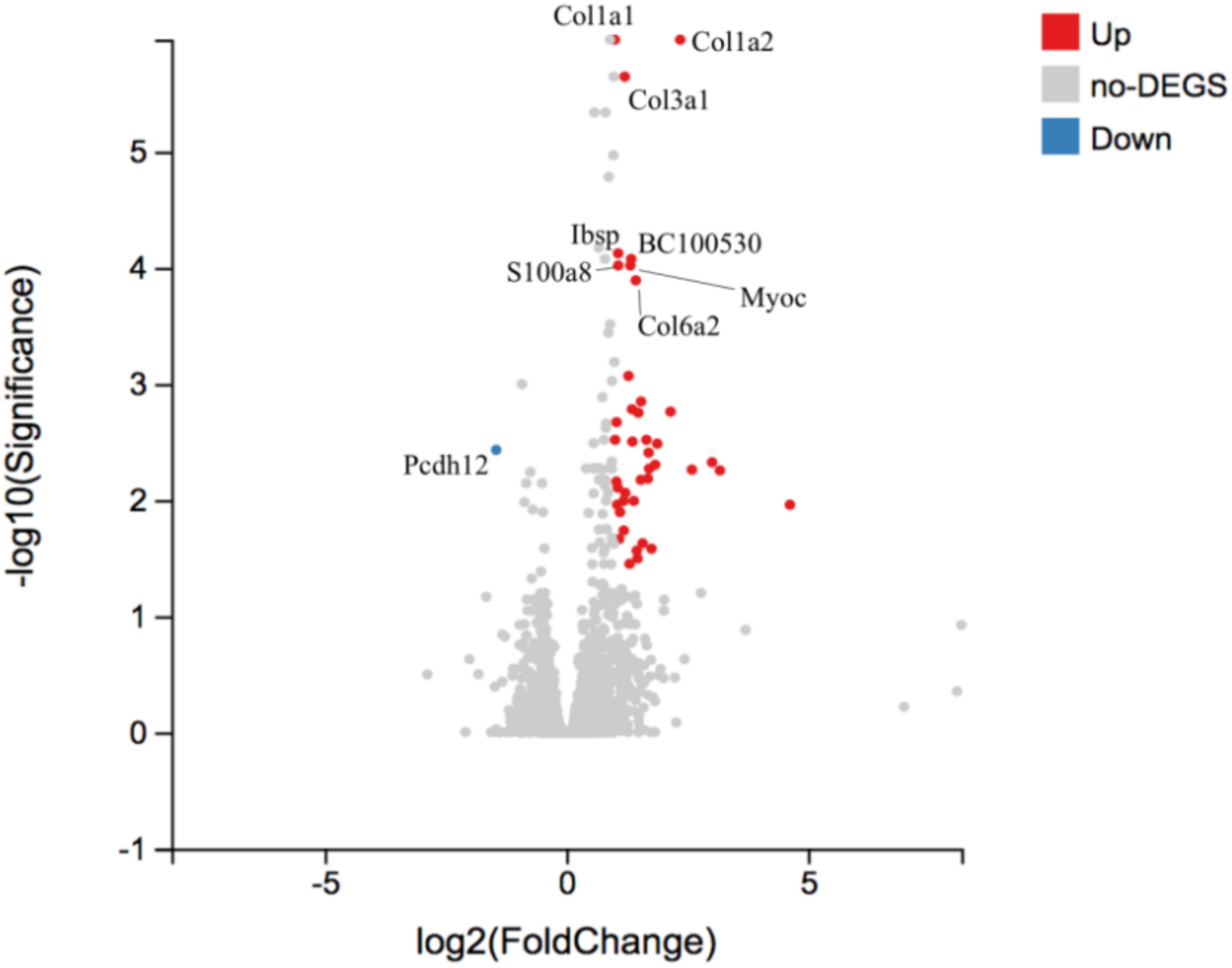
Volcano plot of differentially expressed genes. Up-regulated and down-regulated genes are identified by log2 (Fold Change) with *P* < 0.05. The top 8 up-regulated genes (*Col1a1, Col1a2, Col3a1, Ibsp, BC100530, S100a8, Myoc, and Col6a2*) are represented by red dots, and one significantly down-regulated gene (*Pcdh12*) is represented by the blue dot.

## Discussion

OP is characterized by the imbalance between OC bone resorption and OB bone formation, which leads to bone loss and structural decay, thereby reducing bone strength and increasing fragility. Hence, new therapeutic targets, possibly through the molecular understanding of the communication and command signaling networks between bone formation OB and bone absorption OC, are currently under intense exploration [1].

Recent studies offer new insights into the mechanisms regulating bone and cartilage growth, along with homeostasis [14]. The *Wnt* signaling pathway plays a crucial role in bone development and homeostasis [3]. Both the canonical and non-canonical *Wnt* signaling pathways are activated by the binding of the *Wnt* ligand to *Fzd* receptors, alone or in combination with specific co-receptors [15]. Studies have shown that *FZD8*, which plays a significant role in bone remodeling, is highly expressed in OC and OB [9]. Hence, *Fzd8* might be a potential therapeutic target for OP, although its signal transduction mechanism has not been elucidated [6].

Different variants contributing to heritability tend to spread across the whole genome, which might explain why some loci/genes characterized by genome-wide association studies for BMD are considered not to influence OP pathophysiology, while other genes tend to be core genes [16].

We discovered that in *Fzd8-*knockout mice, *P1NP* decreases, and *CTX* increases. The micro-CT results showed that the trabeculae, BV, and BMD decreased after *Fzd8*-knockout. Statistically significant differences in *Itgb3* and *RANK* protein expressions were observed in the WT and HO mice based on IHC results. Moreover, in vitro and transgenic animal studies have demonstrated that *Itgb3* has important functions in OC resorptive mechanisms [17]. *Fzd10* expression, which positively regulates the *Wnt* signaling pathway, is significantly down-regulated in OP [18]. In this study, the genes *Col1a1*, *Col1a2*, *Col3a1*, *Ibsp*, *S100a8*, *BC100530*, *Myoc*, *Col6a2*, and *Pcdh12* were differentially expressed (*P* < 0.01, **Figure 5**). These genes are potentially relevant to OP [19], which further elucidates the effects of *Fzd8*-knockout on OP.

To date, only a few SNPs/genes and their functional mechanisms have been successfully characterized in OP [20]. By knocking out the *Fzd8* gene, we observed significant differences in the expressions of the *Fzd10, Lta, Itgb3*, and *RANK* proteins in the WT and HO mice. At present, proteins of the frizzled family, including *FZD10*, may act as Wnt co-receptors on the cell membrane [21]. The *FZD* protein activates intracellular signal transduction molecules and regulates the target gene expressions. Moreover, sFRPs could inhibit *Wnt* protein activity in bone pathophysiology regulation, bone mineral content, and immune/inflammatory responses [3].

To identify such receptors, previous studies assessed the expression of known *Fzd* genes in distinguished tissues and bone cells [9]. Moreover, a study found that *Itgb3* is highly expressed in OC, which is the most important integrin regulating OC function [22]. *TNF-α* antibodies can decrease *Itgb3* and V-ATPase expressions in OC. Therefore, *TNF-α* may enhance OC bone resorption by increasing the *Itgb3* and V-ATPase expression in OC [23].

Inflammatory cytokines are highly active and multi-functional small molecule proteins that are mainly generated by immune cells [24]. In this study, with *Fzd8*-knockout, significant differences in *Lta* gene expression were observed among the WT and HO mice. Previous investigations have shown that *Lta* activates immune responses via downstream signaling pathways, including the expression of *NF-κB* [25]. Osteogenic markers such as alkaline phosphatase, *type-I collagen*, and *Runx2* are up-regulated in *Lta*-stimulated mesenchymal stem cells [26]. Our data showed that *Lta* expression in the HO *Fzd8*-knockout mice was significantly lower than that in WT mice. Previous reports have shown that *Lta* could promote the growth of OB indirectly, verifying that *Fzd8*-knockout could promote OP [26].

In contrast, *TNF-α* exerts its function through each of two receptors, i.e., *TNFR1* and *TNFR2*, that either contains [27] or lacks a death domain [28], respectively. *TNFR1* regulates OC formation to function positively [29]. Although *TNF-α* is capable of activating the *NF-κB*, *JNK*, *p38*, *ERK,* and *Akt* pathways in OC precursors and OCs, the signaling cascades leading to pathway activation have not been established for OC precursors [25]. Many inflammatory cytokines, including *IL-6* and *TNF-α,* participate in OC differentiation [30]. Our data show that *Fzd10* is significantly down-regulated, while *Itgb3* and *RANK* are significantly up-regulated in OP. This suggests that the corresponding differentially expressed miRNAs might regulate OP by suppressing positive regulators, which is in agreement with the findings of Albers et al. [9]. In combination with the perspective of WGS data, it is found that *Lta* is significantly down-regulated in OP. As an important factor of inflammatory response, *Lta* may be involved in the immune regulation of *TNF* on OC and OB during OP, but the special mechanisms remain to be elucidated, which could provide us with new insights for further research and another promising direction for precise treatment of OP.

Our *Fzd8*-knockout murine model showed that there were significant alternations in *Fzd10* and *Lta* gene expressions at RNA level and *Itgb3* and *RANK* protein expressions between WT and HO mice, which are significantly associated with bone remodeling. Our results revealed that *FZD8* could be a therapeutic target in OP. This study elucidates the molecular mechanisms in OP, providing evidence-based data for OP drug development and treatment.

## Acknowledgments

This study was funded by the National Natural Science Foundation of China (grant No. 81272168), the Key Clinical Specialty Discipline Construction Program of Fujian, P.R. China and the Health Youth Research Project of Fujian Province (grant No. 2020QNB060). We express gratitude to Chengqi He for providing guidance, and Hesong Qiu for processing the data and managing this project.

## Author contributions

Zhengkun Lin, Jianquan He, and Jian Chen conceived and designed all experiments; Zhengkun Lin, Hui Huang, Jianquan He, Xiaomei Lin, and Heqing Chen performed the experiments; Wen Zhang analyzed the data; Zhengkun Lin, Jianquan He, Wen Zhang and Jian Chen wrote the paper. All authors have read and approved the final manuscript.

## Availability of data and materials

All data generated or analyzed in the study are included either in this article or in the supplementary files. Relevant data are available from the corresponding author, upon request.

## Compliance with Ethical Standards

All animal experiments were carried out in Zhongshan Hospital Xiamen University following the Instructive Notions with Respect to Caring for Laboratory Animals issued by the Ministry of Science and Technology of People’s Republic of China.

## Ethical approval

The study was confirmed and approved by the Animal Subjects Committee of Zhongshan Hospital Xiamen University (approval no. XMVLAC20120044).

## Consent for publication

Not applicable.

